# Effects of combined exposure to ethanol and delta-9-tetrahydrocannabinol during adolescence on synaptic plasticity in the prefrontal cortex of Long Evans rats

**DOI:** 10.1101/2023.08.14.553087

**Authors:** Linyuan Shi, Shuo Kang, Chan Young Choi, Brynn L. Noonan, Lauren K. Carrica, Nu-Chu Liang, Joshua M. Gulley

## Abstract

Significant exposure to alcohol or cannabis during adolescence can induce lasting disruptions of neuronal signaling in brain regions that are later to mature, such as the medial prefrontal cortex (mPFC). Considerably less is known about the effects of alcohol and cannabis co-use, despite its common occurrence. Here, we used male and female Long-Evans rats to investigate the effects of early-life exposure to ethanol, delta-9-tetrahydrocannabinol (THC), or their combination on high frequency stimulation (HFS)-induced plasticity in the prelimbic region of the mPFC. Animals were injected daily from postnatal days 30 to 45 with vehicle or THC (escalating doses, 3-20 mg/kg) and allowed to drink vehicle (0.1% saccharin) or 10% ethanol immediately after each injection. *In vitro* brain slice electrophysiology was then used to record population responses of layer V neurons following HFS in layer II/III after 3-4 weeks of abstinence. We found that THC exposure reduced body weight gains observed in *ad libitum* fed rats, and reduced intake of saccharin and ethanol. Compared to controls, there was a significant reduction in HFS-induced long-term depression (LTD) in rats exposed to either drug alone, and an absence of LTD in rats exposed to the drug combination. Bath application of indiplon or AR-A014418, which enhance GABA_A_ receptor function or inhibit glycogen synthase kinase 3β (GSK3β), respectively, suggested the effects of ethanol, THC or their combination were due in part to lasting adaptations in GABA and GSK3β signaling. These results suggest the potential for long-lasting adaptations in mPFC output following co-exposure to alcohol and THC.

## INTRODUCTION

Alcohol and cannabis are two drugs whose use is widespread and often initiated in adolescence. Estimates from the 2019 National Survey on Drug Use and Health suggest 37% and 57% of the 12-25 year olds in the U.S. had initiated cannabis and alcohol use, respectively (SAMHSA, 2020). Frequent use of alcohol or cannabis has been associated with cognitive impairments in adolescent-onset users who took these drugs regularly and at high doses, or who met diagnostic criteria for substance use disorder (Ehrenreich et al., 1999; Lees et al., 2020; Thoma et al., 2011). The mechanisms underlying these drug use-associated cognitive disruptions are not yet understood, but many studies have suggested that late-maturing brain regions such as the medial prefrontal cortex (mPFC) may be especially susceptible to drug-induced adaptations when exposure occurs in adolescence compared to adulthood (Gulley & Juraska, 2013). The mPFC undergoes significant anatomical and functional alterations throughout peri-adolescence, and this has been suggested to underlie the maturation of higher cognitive functions such as memory, attention, behavioral flexibility and impulse control that characterize the transition to adulthood (Drzewiecki & Juraska, 2020; Euston et al., 2012; Gogtay et al., 2004).

A key feature of mPFC maturation is an increase in the activity of gamma-amino butyric acid (GABA)-releasing fast-spiking interneurons (FSIs), which results in an enhanced inhibitory influence on pyramidal cells in the mPFC (Caballero et al., 2014a; Caballero et al., 2016). Electrophysiology studies suggest that this heightened inhibitory tone may allow for the expression of high frequency stimulation (HFS)-induced long-term depression (LTD) in the mPFC, which emerges around late adolescence to young adulthood. For example, in some rats that were 45-55 days old and all rats that were ≥ 65 days old, HFS (four stimulus trains at 50 Hz) delivered in the ventral hippocampus induced a significant and long-lasting decrease in evoked field potentials in layer 5/6 of the mPFC (Caballero et al., 2014b). This stimulation-induced LTD in the deep layer mPFC is also observed following HFS delivered in layer 2/3 of the mPFC, where it begins to emerge in rats as young as 55 days old (Kang et al., 2018). Notably, early adolescent exposure to delta-9-tetrahydrocannabinol (THC) or WIN 55,212-2, which are agonists at type 1 cannabinoid (CB_1_) receptors, disrupts the ability of the mPFC to undergo HFS-induced LTD in adulthood (Cass et al., 2014; Rubino et al., 2015). The effects of adolescent alcohol exposure on HFS-induced LTD in the mPFC has not, to our knowledge, been reported previously. However, binge-like exposure to ethanol during adolescence has been shown to significantly reduce GABA_A_ receptor-mediated inhibitory tone in the PFC (Centanni et al., 2017; Hughes et al., 2019).

Co-use of alcohol and cannabis may be much more common than using either drug alone, with studies estimating from 30% to 80% of adolescent users of alcohol or cannabis co-use the drugs on at least some occasions (Pape et al., 2009; Subbaraman & Kerr, 2015; Terry-Mcelrath & Patrick, 2018; Yurasek et al., 2017). This co-use, when heavy or associated with a substance use disorder, has been associated with persistent deficits in cognitive function and adverse psychosocial outcomes (Lees et al., 2021). A recent longitudinal imaging study in over 700 adolescents suggested faster declines in gray matter volume in multiple brain regions, including the frontal cortex, in participants that co-used alcohol and cannabis compared to those who used either drug alone (Luo et al., 2022). Published studies of co-exposure in laboratory rodents, which are relatively sparse despite the prevalence of co-use in adolescents, have thus far reported modest or no behavioral effects of co-exposure compared to alcohol or THC exposure alone (Liang, 2022). For example, we recently found that self-administration of alcohol and THC during peri-adolescence had no effect on attention, working memory, or behavioral flexibility when behavior was assessed in young adulthood (Carrica et al., 2023). No unique effects of co-exposure were also reported in studies of recognition memory, instrumental learning, fear conditioning, and spatial memory (Hamidullah et al., 2021; Smiley et al., 2021). However, co-exposure may induce more significant adaptations in PFC function. In a study that used a concurrent exposure approach, where male rats were exposed to vaporized THC from postnatal day (P) 45 to P50 and vaporized ethanol from P50 to P64, there was an enhanced calcium signaling response in the prelimbic cortex to conditioned fear stimuli that was absent in rats exposed to either drug alone (Smiley et al., 2021). Thus, co-exposure to alcohol and cannabis may uniquely impact the function of the mPFC, even when such adaptations are not expressed at the level of behavior.

In the current study, we examined whether peri-adolescent exposure to ethanol, THC or both drugs would alter the expression of HFS-induced plasticity in the mPFC. Rats were allowed to voluntarily drink sweetened ethanol (10% in 0.1% saccharin) and were injected (s.c.) with vehicle or escalating doses of THC (3-20 mg/kg). They were sacrificed during young adulthood, about three weeks after the last drug exposure, for *in vitro* brain slice recordings of population responses in layer 5/6 of the prelimbic cortex. Our hypothesis, which was based on our previous work (Kang et al., 2018) showing rats normally develop the ability to express HFS-induced LTD in the mPFC during the age range when drug exposure occurred, was that drug exposure would reduce the magnitude of LTD and combined exposure would have the greatest impact. To test potential mechanisms underlying drug-induced changes in HFS-induced plasticity, we also performed recordings when indiplon, a positive allosteric modulator of GABA_A_ receptors, or AR-A014418, an inhibitor of glycogen synthase kinase 3β (GSK3β), were applied to the brain slices. Considering previous work showing the effects of peri-adolescent drug exposure on GABA function in the mPFC (Caballero et al., 2014b; Centanni et al., 2017), and that repeated exposure to these drugs may upregulate GSK3β signaling (He et al., 2006), we hypothesized that indiplon and AR-A014418 would reverse the effects of ethanol and THC on HFS-induced LTD.

## MATERIALS AND METHODS

### Animals

We used a total of 64 Long Evans rats (32 male, 32 female) that were ordered from Envigo (Indianapolis, IN, USA) and arrived at our colony at approximately postnatal day (P) 24 (± 2 days). Four cohorts of rats that were received over a period of 7 months were used for these experiments, with two rats from each cohort represented in each of the groups in the study. Rats were housed in a temperature-controlled room on a 12:12 light/dark cycle (lights on at 1000 h) with food and water available *ad libitum.* Rats were pair-housed with same-sex rats and weighed daily from the arrival date to the end of the drug treatment period (P45). They remained pair-housed until they were sacrificed for brain slice electrophysiology between P70 and P88.

Approximately half of the subjects were housed individually for ∼24 h before euthanasia because their cagemate had been removed for sacrifice on the previous day. All experiments were pre-approved by the Institutional Animal Care and Use Committee at the University of Illinois at Urbana-Champaign and were consistent with the Guide for the Care and Use of Laboratory Animals.

### Drugs and exposure procedures

Using our previously described approach (Nelson et al., 2019a), rats were first habituated to the daily procedure of s.c. injection and sipper tube drinking from P28-P29. During the last 1.5 h of the dark cycle and the first 1 h of the light cycle (0830-1100 h), they were injected with 0.15 mL of the THC vehicle and were immediately provided access to sipper tubes containing a 0.1% saccharin solution (saccharin sodium salt hydrate; Sigma-Aldrich). This was done in their homecage, with a clear acrylic divider separating cagemates so their individual consumption of fluid could be monitored. Cagemates remained separated in their home cage for the remainder of the experiment, but the divider contained small holes to allow for potential olfactory and tactile interaction between cagemates.

Following two days of saccharin drinking, rats were pseudorandomly assigned to one of four groups so that total intake was approximately equal between groups (see Supplementary Materials, Table 1). From P30-P45, rats in the control group (CTL) received daily vehicle injections and drank 0.1% saccharin, rats in the ethanol only group (EtOH) were injected with vehicle and drank 10% ethanol (Decon Laboratories) mixed with 0.1% saccharin, rats in the THC only group (THC) were injected with THC and drank 0.1% saccharin, and rats in the combination group (COM) were injected with THC and drank the ethanol-saccharin solution.

The 2.5 h/day drinking paradigm with 10% sweetened ethanol was designed to model low to moderate ethanol drinking behavior (Nelson et al., 2016). THC (200 mg/mL dissolved in 95% ethanol; Research Triangle Institute, distributed by The National Institute on Drug Abuse) was diluted in ethanol (190 Proof; Decon Laboratories), Tween-20 (Sigma-Aldrich), and 0.9% saline in a ratio of 1:1:18 for s.c. injection such that the volume of THC and ethanol together accounted for 1/20 of the total volume of the injection.

The timeline for treatments and sacrifice for brain slice electrophysiology is summarized in Fig. 1. For rats in the THC and COM groups, THC dose was increased from 3 mg/kg on P30-P33, 5 mg/kg on P34-P37, 10 mg/kg on P38-P41, and 20 mg/kg on P42-P45. The highest dose, which was split into two injections to minimize sedation or hypothermia (Taffe et al., 2015), was given as 10 mg/kg at the same time of day as all other vehicle or THC doses (0830) and 10 mg/kg given at ∼2130 h.

**Figure 1.**
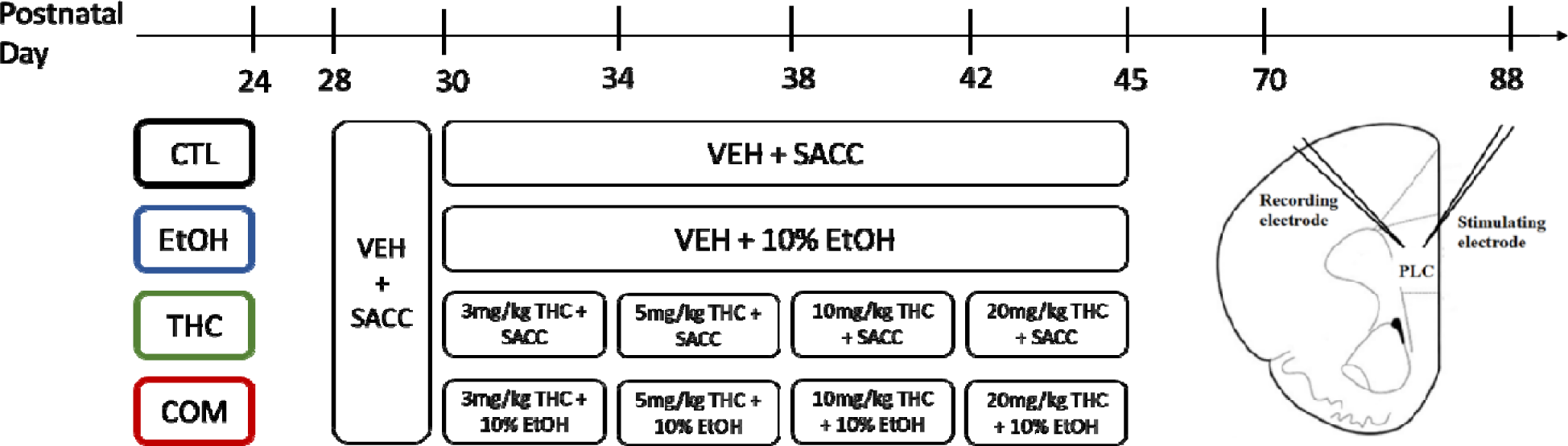
Timeline of treatment and slice electrophysiology. The four groups (*N* = 7-8 rats/sex/group) were control (CTL), which were injected (s.c.) with vehicle and drank 0.1% saccharin (SACC), ethanol only (EtOH), which were injected with vehicle and drank 10% ethanol in SACC, THC only (THC), which were injected with THC and drank SACC, and combination (COM), which were injected with THC and drank 10% ethanol in SACC. Rat brains were harvested for slice electrophysiology between P70 and P88. The schematic diagram shows one hemisphere of a coronal slice and the approximate position of stimulating and recording electrodes (layers II-III and layers V-VI, respectively) in the prelimbic cortex (PLC).

### Brain slice electrophysiology

Between P70 and P88, rats were given an overdose of sodium pentobarbital, perfused transcardially with slicing medium, and decapitated. The brain was immediately removed and put in ice-cold, oxygenated slicing medium containing (in mM): 2.50 KCl, 1.25 NaH2PO4, 10.0 MgCl2, 0.50 CaCl2, 26.0 NaHCO3, 11.0 glucose, and 234 sucrose. Coronal brain slices (450 µm thickness) containing the prelimbic cortex (PLC) region of the medial PFC were obtained with a vibrating tissue slicer (Pelco EasiSlicer, Ted Pella INC.), with the slices subsequently transferred to artificial cerebrospinal fluid (aCSF) containing (in mM): 126 NaCl, 2.50 KCl, 26.0 NaHCO3, 2.00 CaCl2, 1.25 MgCl2, 1.25 NaH2PO4, and 10.0 glucose. Slices were kept in the aCSF oxygenated with 95% O_2_/5% CO_2_ first at 31LJ°C for 20 min and then room temperature for at least 1 h before recordings. During recording sessions, the brain slice on top of the recording chamber (Harvard Apparatus) was perfused in 31LJ°C, oxygenated aCSF while the rest of the slices sit in the room temperature, oxygenated aCSF. A tungsten microelectrode was used as the stimulating electrode and a concentric bipolar microelectrode (Platinum-Iridium inner pole and a Stainless Steel outer pole) as the recording electrode (FHC, Inc., Bowdoin, ME). Recorded signals were amplified (1,000X), filtered (low pass = 10 kHz, high pass = 1 Hz), sampled at 20 kHz and digitally stored.

Using an approach we described previously (Kang et al., 2018), recordings of fEPSPs evoked by a single test pulse (100 µs duration, 190-350 pA at 0.067 Hz) were obtained in layer V/VI of the prelimbic cortex before and after delivery of HFS (4 trains of 100 pulses at 50 Hz/train; 10-s inter-train interval) in layer II/III of the PLC. In the first group of recordings, slices were perfused with aCSF and a 10-min period of baseline fEPSP recordings was obtained with test pulses applied once/min. This was followed by HFS and an additional 40 min of recording following application of test pulses once/min. In a separate group of recordings, drugs (or their vehicle) were added to the aCSF perfusion to determine the effects of pharmacological manipulation of GABA_A_ or GSK-3β signaling of HFS-induced plasticity. The GABA_A_ receptor positive allosteric modulator indiplon (AdooQ Biosciences) or the GSK3β inhibitor AR-A104418 (AdooQ Biosciences) were dissolved in dimethyl sulfoxide (DMSO; Fisher Scientific) as stock solutions (5-10 mM) and kept at −20LJ°C before use. Stock solutions were diluted in 200LJml aCSF oxygenated with 95% O2/5% CO2 (30–31 °C) to make 5 µM Indiplon and 10 µM AR-A014418 solutions in 0.1-0.2% DMSO. When DMSO (control condition), indiplon, or AR-A014418 were perfused, 10 min of baseline activity was recorded followed by 10 min of baseline recording with the compound added to the perfusion medium. This was followed by application of HFS and 40 min of fEPSP recordings with test pulses applied once per min. The DMSO vehicle or drug remained in the perfusion medium for the entirety of the post-HFS recording.

### Data analysis

All statistical analyses were conducted in R using the open-source edition of R Studio (Version 1.4.1717; rstudio.com). Body weight data were analyzed using two-way ANOVA (treatment x day) for each sex separately due to the large body weight differences between females and males under baseline conditions. During-treatment (P30-45) body weight data were presented and analyzed separately from the post-treatment data (measured on P52, P55, P59, P62, and P69) to examine acute and long-term effects of ethanol and THC exposure (see Supplementary Figure 1). Intake of the saccharin and ethanol solutions was measured by subtracting the weight of the sipper tube (to the nearest 0.1g) at the end of each drinking session from the weight before the session. These data were analyzed using three-way ANOVA (sex x treatment x THC dose) followed by two-way ANOVA for each sex separately.

For electrophysiology data, the slope of the first 1-2 ms of the fEPSP was obtained using Clampfit software (version 10.6; Molecular Devices) and averaged in 1-min bins. For each slice, the mean slope was then normalized to the mean slope of its 10-min baseline. Statistical tests consisted of one-way ANOVAs for each treatment group (with four drug perfusion conditions), two-way ANOVAs (sex x treatment) or three-way (sex x treatment x time) ANOVAs. Tukey post-hoc tests were performed when appropriate for all analyses.

## RESULTS

### Effects of drug exposure on body weight and intake of saccharin or ethanol

As expected for free-fed rats of the ages used here, there were significant and rapid increases in body weight from P24 through P68. In addition, there were significant effects of drug exposure on body weight that depended on sex and drug (Fig. 2). Exposure to THC alone or in combination with ethanol ingestion led to a significant decrease in body weight gain in both male and female rats, whereas ethanol intake alone significantly reduced body weight only in females. Two-way ANOVA revealed main effects of treatment (males: F_3, 729_ = 12.1, *p* < 0.001; females F_3, 756_ = 19.4, *p* < 0.001) and day (males: F_26, 729_ = 226, *p* < 0.001; females: F_26, 756_ = 394, *p* < 0.001). No significant interaction was found in either sex. Post-hoc analysis revealed that for males, THC-treated groups (THC and COM) had significantly lower body weights compared to their controls and the ethanol-treated group (Fig. 2A). For females, all drug-treated groups had significantly lower body weights compared to controls, but none differed from each other (Fig. 2B). Further analysis of weight data during and after drug treatment is provided in Supplementary Materials, Fig. 1.

**Figure 2.**
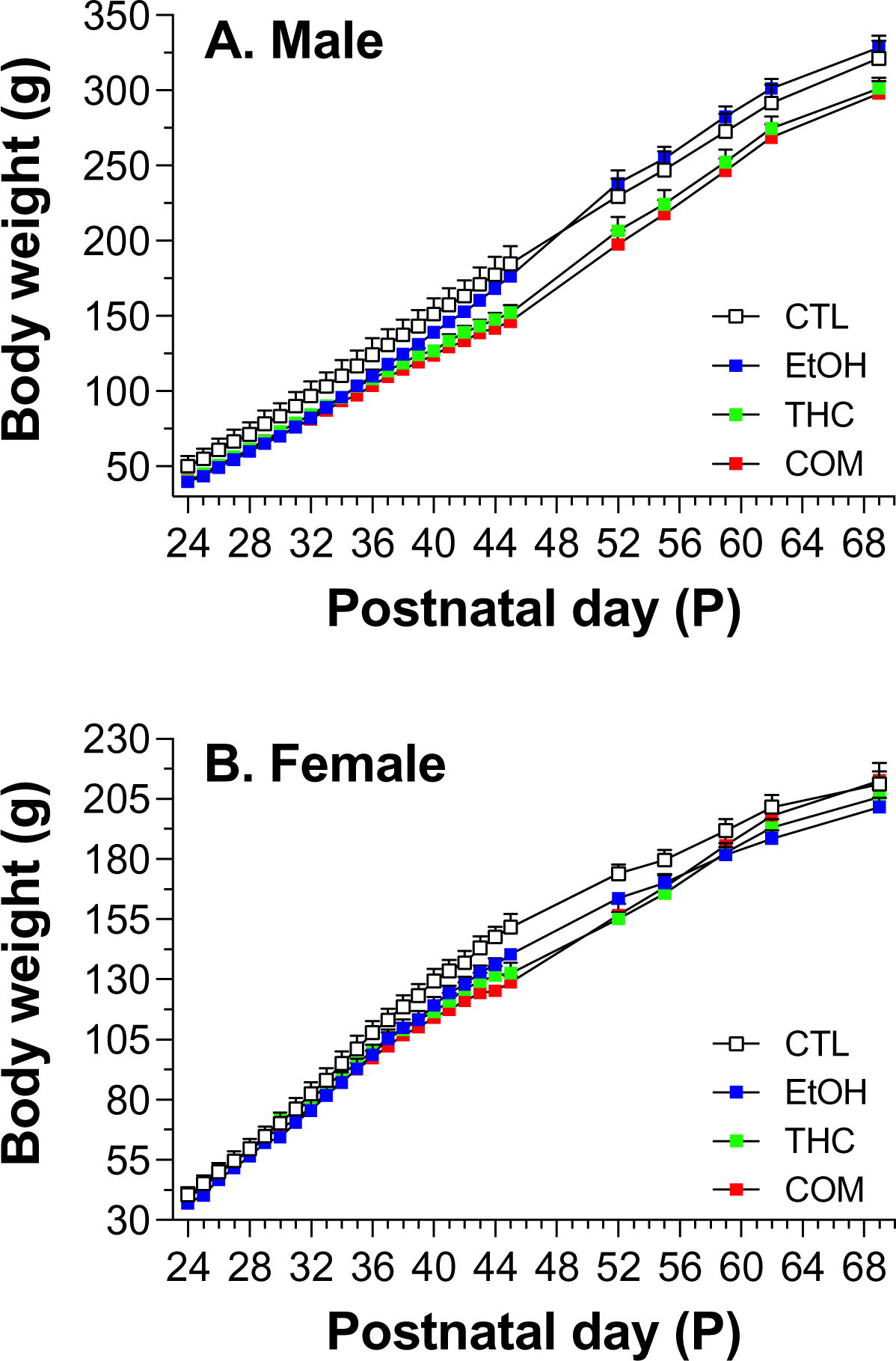
Effects of ethanol, THC, or their combination on body weight of males and females. Weights were measured daily P24-P45, and on P52, P55, P59, P62, and P69. Groups are as described in the legend for Fig. 1 (n = 7-8 rats/sex/group). Two-way ANOVA revealed main effects of treatment (males: F_3, 729_ = 12.1, *p* < 0.001; females F_3, 756_ = 19.4, *p* < 0.001) and day (males: F_26, 729_ = 226, *p* < 0.001; females: F_26, 756_ = 394, *p* < 0.001); there were no significant interactions.

Analysis of saccharin or ethanol intake revealed significant effects of THC on consumption in both males and females (Fig. 3). For saccharin, which was consumed by animals in the CTL and THC groups, a three-way mixed ANOVA (between-subject variables: Sex, Group; within-subject variable: THC dose) revealed main effects of sex (F_1,108_ = 10.7, *p* < 0.001) and treatment (F_1,108_ = 96.8, *p* < 0.001), and significant interactions of sex x treatment (F_1,108_ = 6.65, *p* = 0.011) and treatment x THC dose (F_3,108_ = 7.80, *p* < 0.001). Follow-up two-way mixed ANOVA for each sex (between-subject variables: Group; within-subject variable: THC dose) revealed a significant main effect of treatment for both sexes (males: F_1,52_ = 26.9, *p* < 0.001; females: F_1,56_ = 73.4, *p* < 0.001) and a significant treatment x THC dose interaction for females (F_3,56_ = 5.52, *p* = 0.002). As shown in Fig. 3B, THC exposure suppressed the gradual increase in saccharin intake that was observed in female controls as the treatment period progressed. A similar trend was observed in control males (Fig. 3A), but their increase in saccharin intake over time was lower and the treatment x THC dose interaction was not significant. For ethanol, which was consumed by animals in the EtOH and COM groups, a three-way mixed ANOVA revealed only a main effect of treatment (F_1,112_ = 73.8, *p* < 0.001). Follow-up two-way mixed ANOVA for each sex revealed a significant main effect of treatment for both sexes (males: F_1,56_ = 42.4, *p* < 0.001; females: F_1,56_ = 31.5, *p* < 0.001). As shown in Figs. 3C-D, THC significantly reduced ethanol intake in both sexes, regardless of dose.

**Figure 3.**
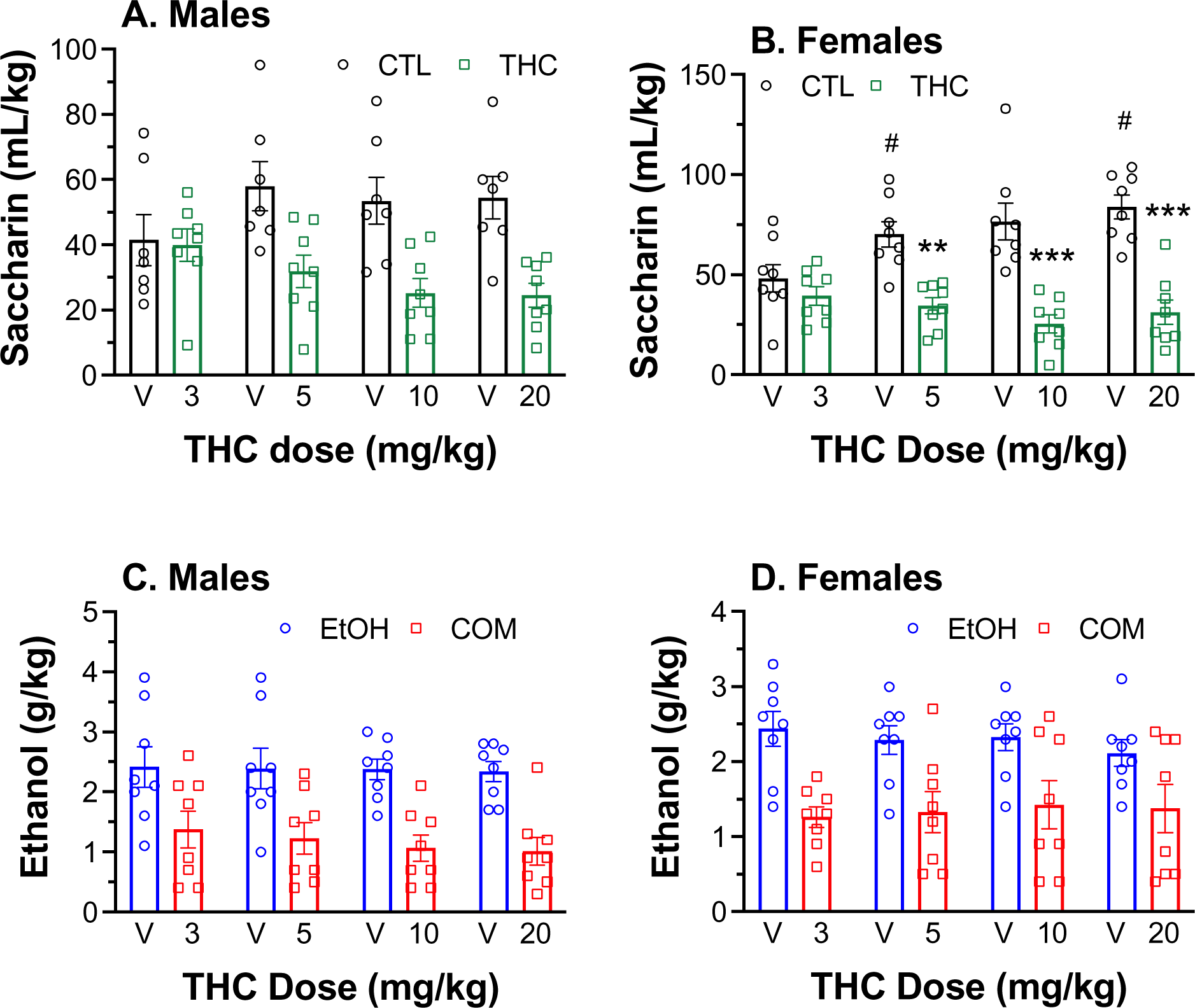
Effects of vehicle (V) or THC injection on the intake of 0.1% saccharin (SACC) or 10% ethanol/SACC. THC doses were escalated across the treatment period (from P30-P45), with intake data presented as mean consumption during the four-day period for the indicated THC dose (or vehicle for rats in the CTL or EtOH groups). Intake for the 20 mg/kg THC dose was measured after the first of the twice daily 10 mg/kg doses. Groups are as described in the legend for Fig. 1 (n = 7-8 rats/sex/group). Two-way ANOVA revealed significant main effect of treatment in A, B, C, and D (all *p* < 0.001), with significant treatment x THC dose interaction in B (*p* = 0.002). \*\**p* < 0.01 and \*\*\**p* < 0.001, compared to CTL; ^#^*p* < 0.05 and ^##^*p* < 0.01, compared to 3 mg/kg dose for THC group or vehicle for CTL group.

### Effects of drug exposure on HFS-induced plasticity

Our initial analysis of the time course of HFS-induced changes in fEPSP slope using three-way ANOVA (sex x treatment x time) revealed significant main effects of sex (F_1,1284_ = 4.07, *p* = 0.044), treatment (F_3,1284_ = 109, *p* < 0.001), time (F_49,1284_ = 12.5, *p* < 0.001), and a sex xtreatment interaction (F_3,1284_ = 5.67, *p* < 0.001). However, post-hoc analyses of the interaction revealed no sex differences within treatment groups. Instead, this interaction was driven by sex differences between different drug treatment groups (e.g., male CTL vs. female COM). For this reason, data were subsequently analyzed and presented collapsed across sex. Presentation of fEPSP data separated by sex is provided in Supplementary Materials, Fig. 2.

As shown in the sample traces (Fig. 4A) and the time course of population response (Fig. 4B), HFS induced a significant reduction in the fEPSP slope in controls. This effect was reduced in drug-treated groups (Fig. 4B), as confirmed by significant main effects of treatment (F_3,1484_ = 116, *p* < 0.001) and time (F_49,1484_ = 13.4, *p* < 0.001). To facilitate treatment group comparisons as in our previous work (Kang et al., 2018), the normalized fEPSP slopes measured during the last 10 min of recordings (30-40 min post-HFS) were used to assess the long-term stable response to HFS (Fig. 4C). Two-way ANOVA of these data revealed a main effect of treatment (F_3,26_ = 7.45, *p* < 0.001), but no effect of sex or interaction. Thus, these data are presented collapsed across sex and were analyzed using one-way ANOVA for treatment effects. We found a significant effect of treatment (F_3,30_ = 7.95, *p* < 0.001), with post-hoc analyses confirming the EtOH and COM groups were significantly different from controls (Fig. 4C). The THC group also exhibited less of an HFS-induced decrease in fEPSP slope compared to controls, but this difference was not statistically significant. The fEPSP slope measured in slices from rats in the COM group showed the least change from baseline following HFS and was significantly different from both the CTL and THC groups.

**Figure 4.**
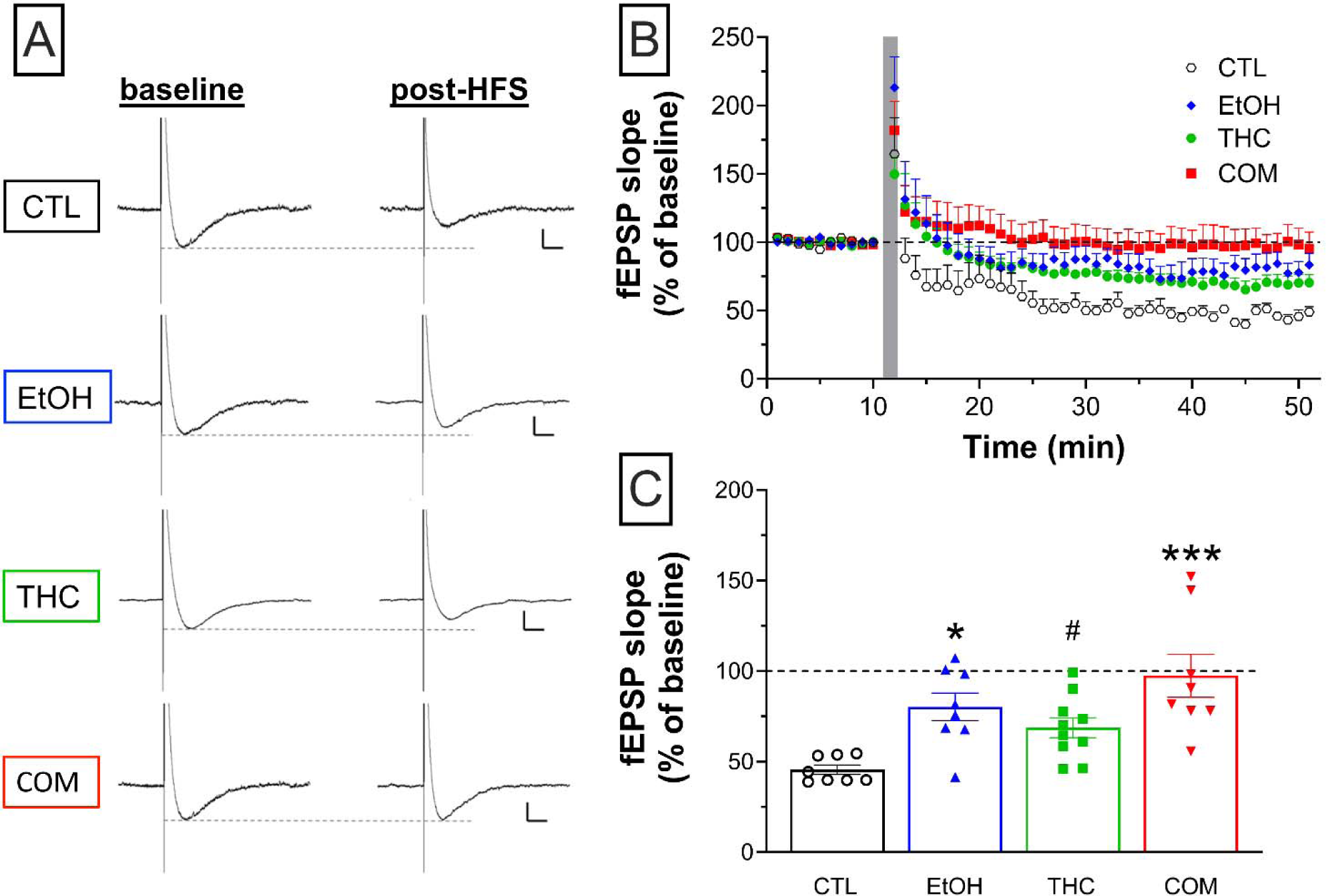
Effects of HFS delivered in layer II/III of the PL on fEPSPs recorded in Layer V/VI. (A) Representative traces sampled before and after application of HFS (4 trains of 100 pulses at 50 Hz/train; 10-s inter-train interval), when brain slices are perfused with aCSF. The vertical line represents the onset of the test pulse (100 μsec duration; 50–350 pA at 0.067 Hz; calibration axes: 500 μV for the ordinate; 100 msec for the abscissa). (B) Time course of the normalized fEPSP slope before and after HFS (shaded vertical bar). Data are presented in 1-min bins. (C) Long-term stable response to HFS. Plotted are the mean fEPSP slopes for the 30-40 min period post-HFS. Groups are as described in the legend for Fig. 1, and data points for individual slices are shown in panel C. \**p* < 0.05 and \*\*\**p* < 0.001 vs. CTL; ^#^*p* < 0.05 vs. COM; n = 8-10 slices from 6-8 rats/group

### Effects of DMSO, indiplon and AR-A104418 on HFS-induced plasticity

In a separate set of recordings, brain slices were perfused for 10 min with the drug perfusion vehicle (dimethyl sulfoxide, or DMSO), indiplon (GABA_A_ positive allosteric modulator) or AR-A014418 (GSK3-β inhibitor). This occurred after a 10-min baseline period (aCSF only) in one of three conditions: 1) 0.1-0.2% DMSO only (shortened as “DMSO” onward); 2) 5 µM indiplon in 0.1-0.2% DMSO (shortened as “indiplon”); or 3) 10 µM AR-A014418 in 0.1-0.2% DMSO (shortened as “AR”). The recordings obtained prior to HFS delivery with aCSF alone or aCSF plus DMSO, indiplon or AR-A014418 were similarly stable and unchanged in all treatment groups, as shown in the representative traces during baseline for brain slices from rats in the different treatment groups (panel A in Figs. 5-8). The descriptive statistics for these data are included in Supplementary Materials, Table 2. Following HFS, there were significant effects of the perfusion compound on the HFS-induced LTP response observed under standard recording conditions (i.e., aCSF perfusion) that depended on the pre-treatment group. For slices from rats in the CTL group (Fig. 5), perfusion of DMSO, indiplon and AR-A014418 led to more variable responses to HFS such that the post-HFS reduction in fEPSP slope was attenuated. This effect was most pronounced following AR-A014418 perfusion, with the fEPSP slope increasing or not changing from baseline in three of the eight recordings at 30-40 min post-HFS (Fig. 5C). One-way ANOVA of these data revealed a significant effect of perfusion (F_3, 26_ = 3.19, *p* = 0.040), and post-hoc tests confirmed that AR-A014418 attenuated HFS-induced LTD compared to the aCSF perfusion group (Fig. 5C).

**Figure 5.**
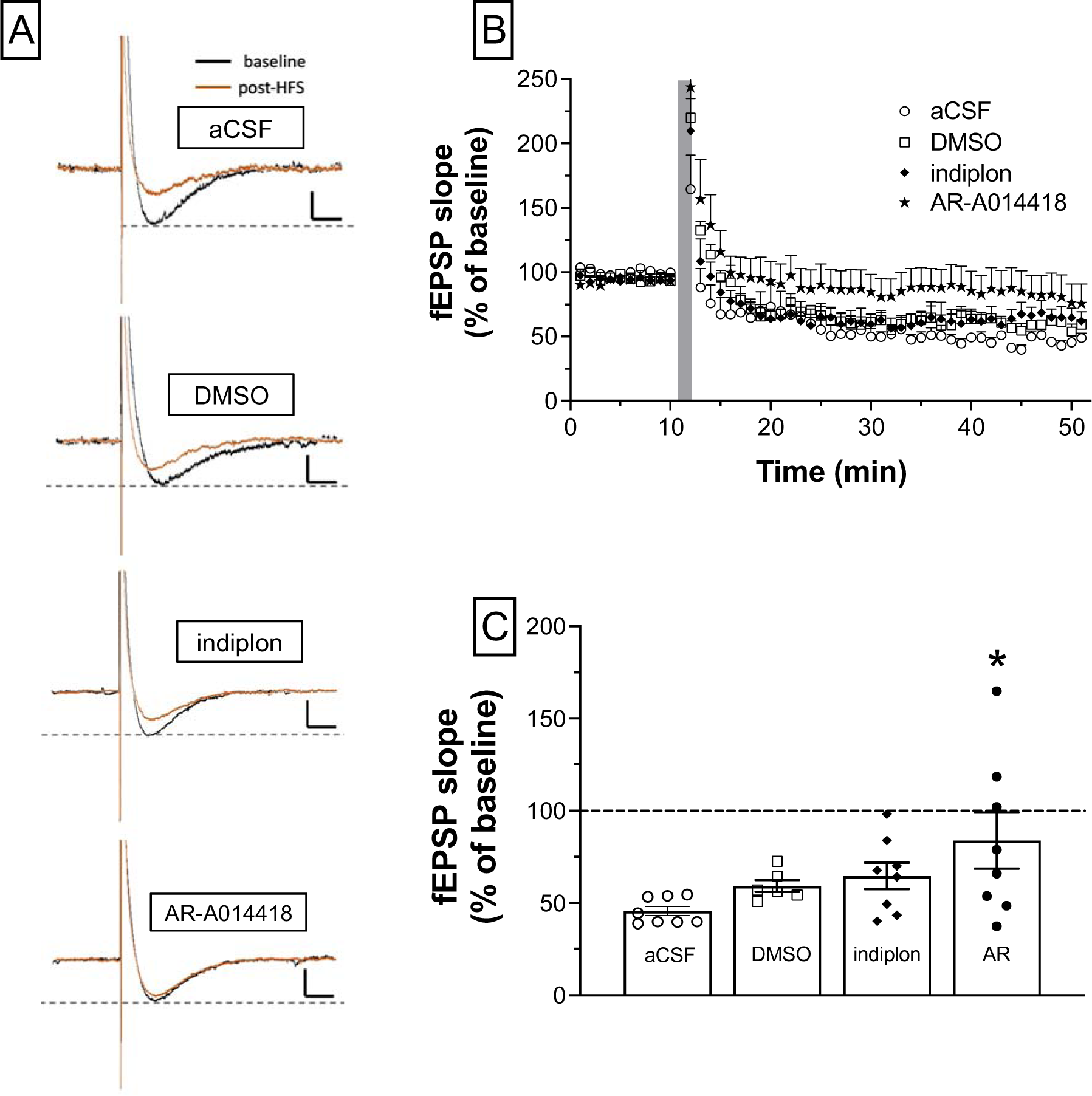
Effects of 0.1-0.2% DMSO, 5 μM indiplon, and 10 μM AR-A014418 on HFS-induced changes in the slope of the fEPSP in brain slices from rats in the control (CTL) group. A) Representative traces sampled before and after application of HFS in all four perfusion conditions (black line: pre-HFS; red line: post-HFS). The vertical line represents the onset of the HFS (100 μsec duration; 50–350 pA at 0.067 Hz; calibration axes: 500 μV for the ordinate; 100 msec for the abscissa). (B) Time course of the normalized fEPSP slope before and after HFS (shaded vertical bar). Data are presented in 1-min bins. C) Long-term stable response to HFS. Plotted are the mean fEPSP slopes for the 30-40 min period post-HFS. Data for the aCSF condition are the same as shown for the CTL group in Fig. 4. Data points for individual slices are shown in panel C; \**p* < 0.05 vs. aCSF; n = 6-8 slices from 6-8 rats/group.

**Figure 6.**
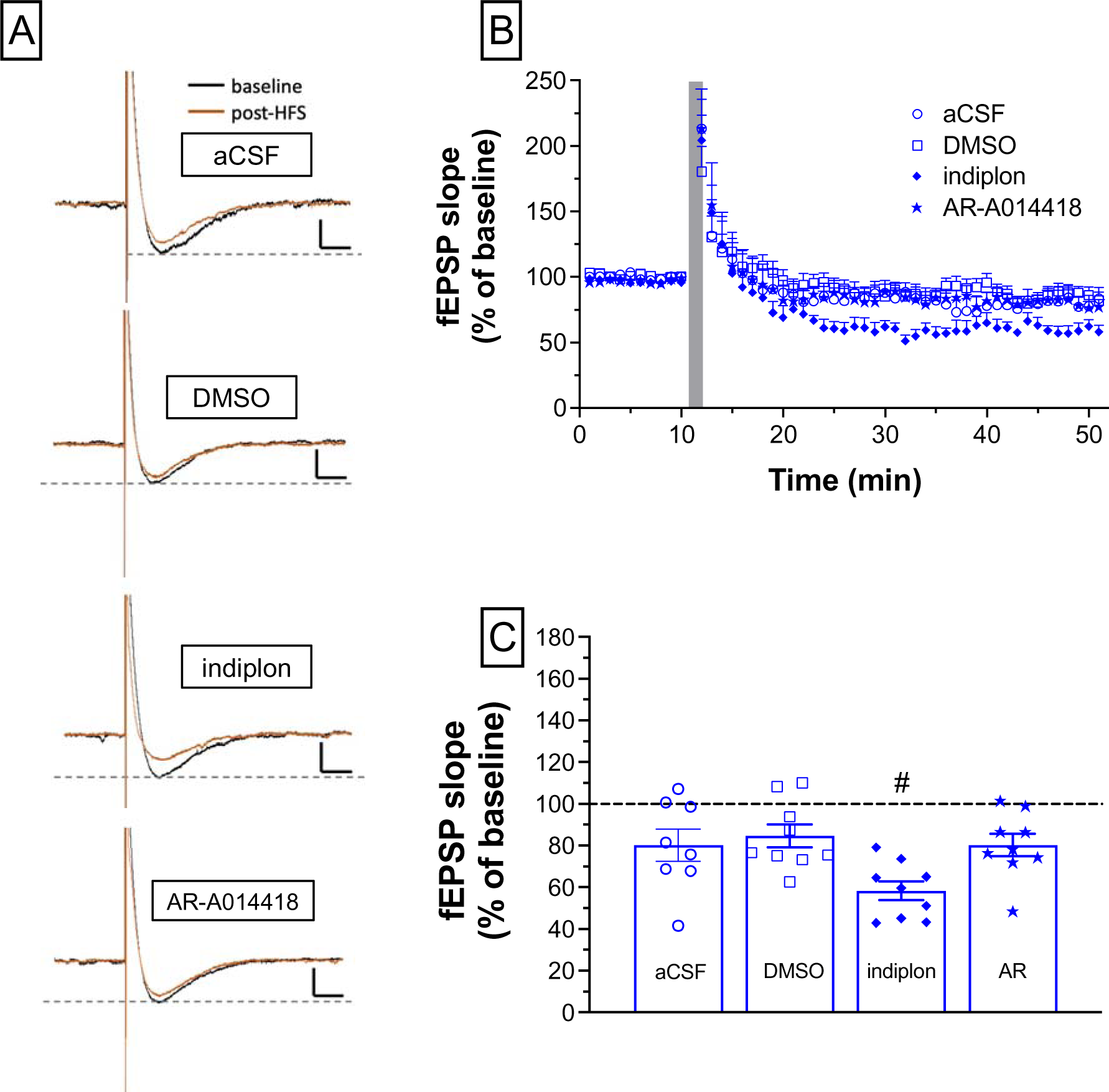
Effects of 0.1-0.2% DMSO, 5 μM indiplon, and 10 μM AR-A014418 on HFS-induced changes in the slope of the fEPSP in brain slices from rats in the ethanol drinking (EtOH) group. A) Representative traces sampled before and after application of HFS in all four perfusion conditions (black line: pre-HFS; red line: post-HFS). The vertical line represents the onset of the HFS (100 μsec duration; 50–350 pA at 0.067 Hz; calibration axes: 500 μV for the ordinate; 100 msec for the abscissa). (B) Time course of the normalized fEPSP slope before and after HFS (shaded vertical bar). Data are presented in 1-min bins. C) Long-term stable response to HFS. Plotted are the mean fEPSP slopes for the 30-40 min period post-HFS. Data for the aCSF condition are the same as shown for the EtOH group in Fig. 4. Data points for individual slices are shown in panel C; ^#^*p* < 0.05 vs. DMSO; n = 8-9 slices from 8-9 rats/group.

**Figure 7.**
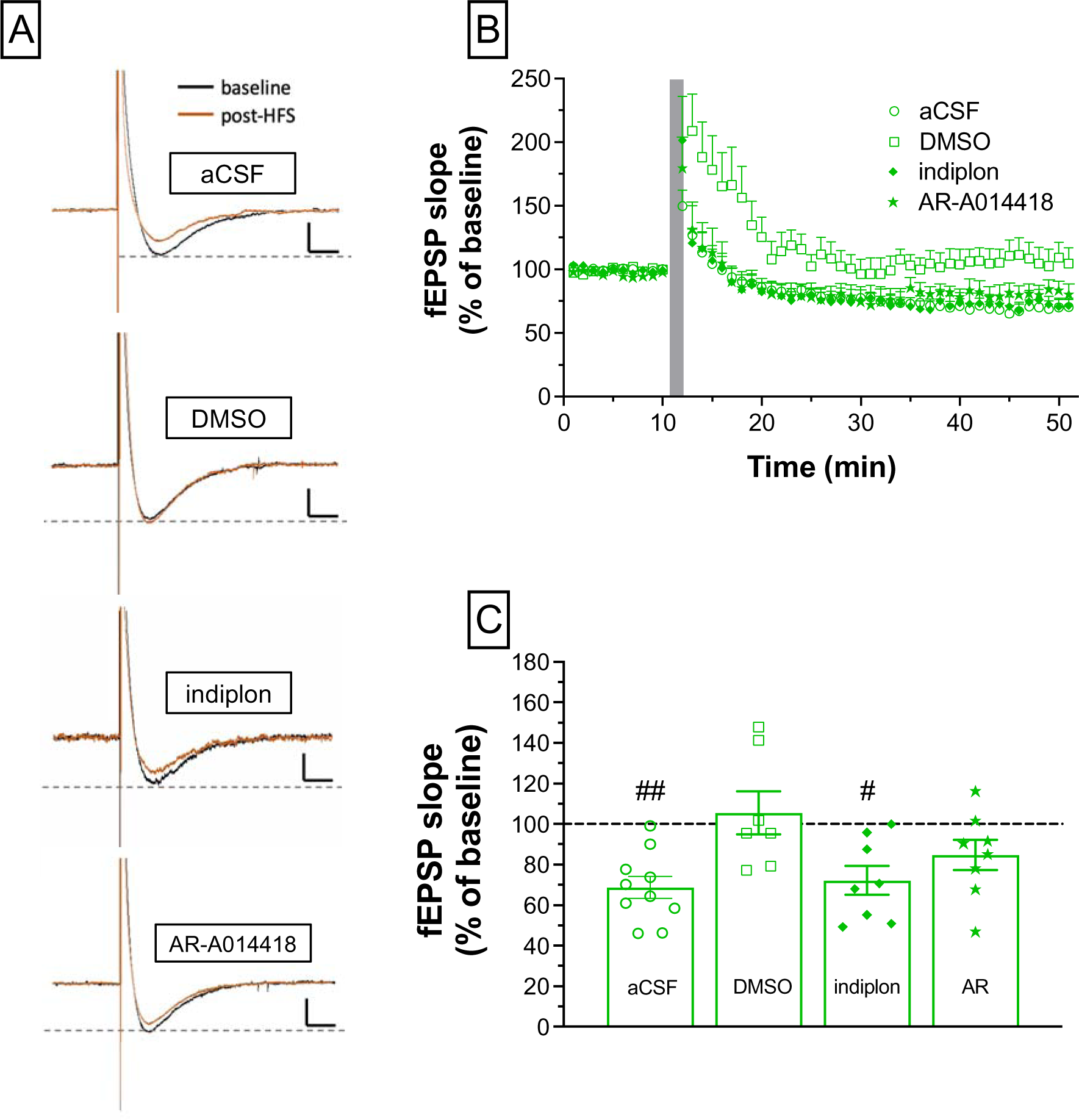
Effects of 0.1-0.2% DMSO, 5 μM indiplon, and 10 μM AR-A014418 on HFS-induced changes in the slope of the fEPSP in brain slices from rats in the THC-exposed group. A) Representative traces sampled before and after application of HFS in all four perfusion conditions (black line: pre-HFS; red line: post-HFS). The vertical line represents the onset of the HFS (100 μsec duration; 50–350 pA at 0.067 Hz; calibration axes: 500 μV for the ordinate; 100 msec for the abscissa). (B) Time course of the normalized fEPSP slope before and after HFS (shaded vertical bar). Data are presented in 1-min bins. C) Long-term stable response to HFS. Plotted are the mean fEPSP slopes for the 30-40 min period post-HFS. Data for the aCSF condition are the same as shown for the THC group in Fig. 4. Data points for individual slices are shown in panel C; \*\**p* < 0.01 vs. aCSF; ^#^*p* < 0.05 vs. DMSO; *n* = 7-10 slices from 6-8 rats/group.

**Figure 8.**
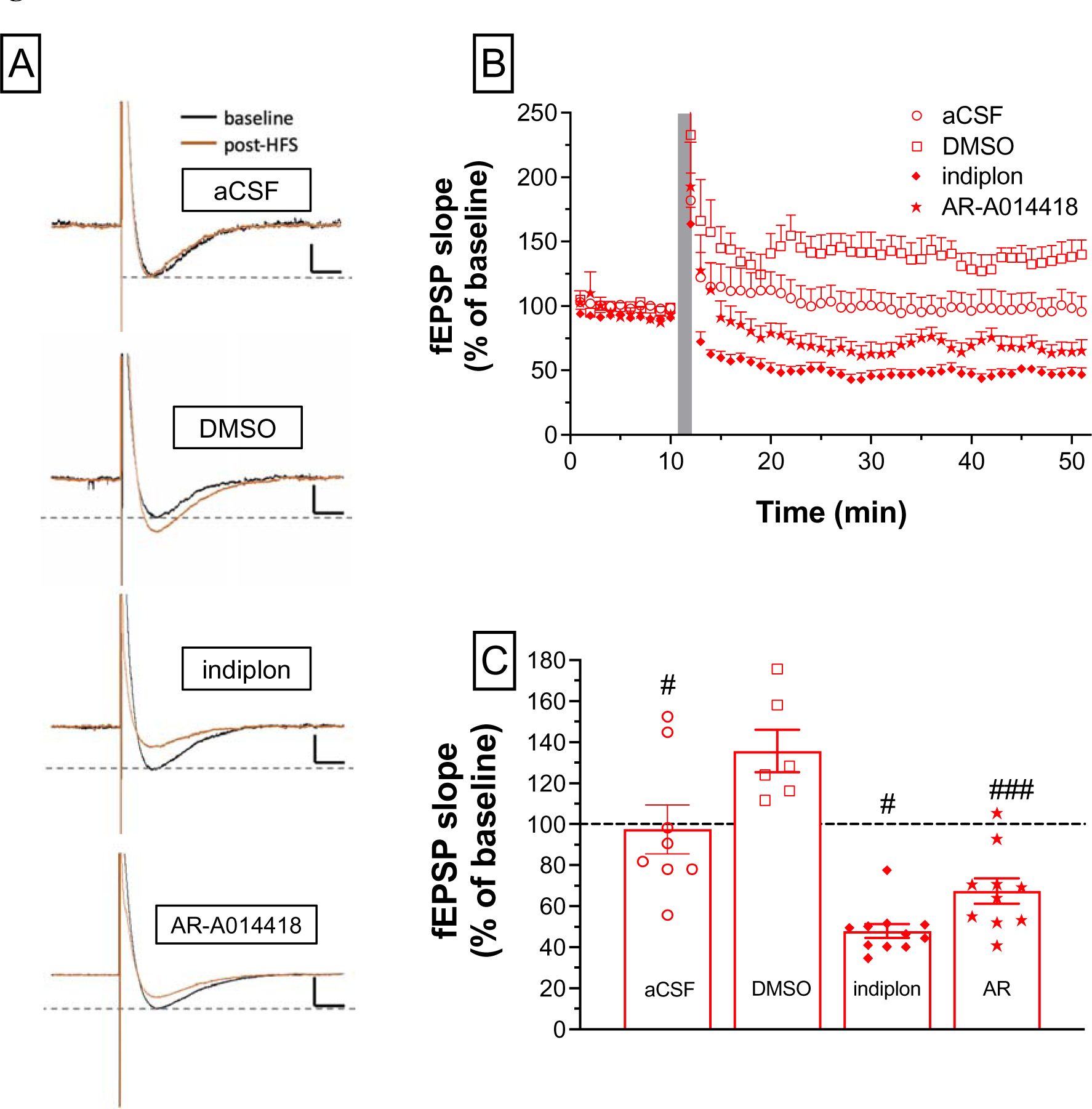
Effects of 0.1-0.2% DMSO, 5 μM indiplon, and 10 μM AR-A014418 on HFS-induced changes in the slope of the fEPSP in brain slices from rats that drank ethanol and were exposed to THC (COM). A) Representative traces sampled before and after application of HFS in all four perfusion conditions (black line: pre-HFS; red line: post-HFS). The vertical line represents the onset of the HFS (100 μsec duration; 50–350 pA at 0.067 Hz; calibration axes: 500 μV for the ordinate; 100 msec for the abscissa). (B) Time course of the normalized fEPSP slope before and after HFS (shaded vertical bar). Data are presented in 1-min bins. C) Long-term stable response to HFS. Plotted are the mean fEPSP slopes for the 30-40 min period post-HFS. Data for the aCSF condition are the same as shown for the COM group in Fig. 4. Data points for individual slices are shown in panel C; \**p* < 0.05 and \*\*\**p* < 0.001 vs. aCSF; ^#^*p* < 0.05 and ^###^*p* < 0.001 vs. DMSO; *n* = 6-11 slices from 6-11 rats/group.

In brain slices from rats in the ethanol drinking group (Fig. 6), the significant reduction in HFS-induced LTD that was observed during standard recording conditions was not altered significantly by perfusion of DMSO or AR-A014418. However, perfusion of indiplon restored HFS-induced LTD to the levels observed in rats from the CTL group (∼40% and ∼45% of baseline, respectively). One-way ANOVA of these data revealed a significant effect of perfusion (F_3, 31_ = 4.44, *p* = 0.011), and subsequent post-hoc analysis confirmed a significant difference in the stable, post-HFS change in fEPSP slope in slices exposed to indiplon compared to its DMSO vehicle (Fig. 6C). In rats exposed to THC (Fig. 7), there was a significant effect of perfusion (F_3, 29_ = 4.63, *p* = 0.009) that was driven by the lack of HFS-induced LTD in slices perfused with DMSO (Fig. 7C). In slices from rats exposed to both ethanol and THC, which was the pre-treatment group with the largest reduction in HFS-induced LTD during aCSF perfusion (∼ 97% and ∼ 45% of baseline for the COM and CTL groups, respectively), we observed a similar effect of DMSO but with the stable response changing to an LTP-like response (Fig. 8). We found a significant effect of perfusion (F_3, 31_ = 22.0, *p* < 0.001), and post-hoc tests revealed both indiplon and AR-A014418 partially restored HFS-induced LTD in these slices (Fig. 8C).

## DISCUSSION

A primary goal of this study was to determine if peri-adolescent exposure to ethanol, THC or both drugs would alter the expression of HFS-induced plasticity in the mPFC. Using *in vitro* extracellular recordings of fEPSPs, we demonstrated that chronic exposure to THC and/or ethanol during peri-adolescence led to a long-lasting decrease in the magnitude of LTD in the deep layers (V/VI) of the mPFC in response to HFS stimulation in superficial layers (II/III). This impaired capacity to undergo plasticity was most evident in rats exposed to both drugs, followed by rats receiving ethanol alone and THC alone. Indiplon, a positive allosteric modulator of the GABA_A_ receptor, partially restored LTD in rats exposed to ethanol alone, THC alone, or both drugs. In the combined-exposure group, we also found evidence for the involvement of the GSK3β signaling pathway in the disruptive effects of drug exposure on HFS-induced plasticity in the mPFC. The selective inhibitor of GSK3β, AR-A014418, partially restored LTD in brain slices from rats exposed to both ethanol and THC. Taken together, these data reveal potential mechanisms through which alcohol and THC act separately and together to exert a persistent influence on the function of the mPFC.

Our body weight change data suggested that daily subcutaneous THC injection led to suppression of weight gain during the treatment period, whether it was administered alone or in combination with alcohol. The combined exposure seemed to extend this effect in female rats post-treatment (at least until P69), but not males, and THC injection alone did not have extending effects on body weight increase after the drug was cleared from the body. This effect of THC is likely due to enhanced metabolism, more specifically improved insulin sensitivity and decreased fat deposition (Nelson et al., 2019b). The significant effects of drug exposure on ingestive behavior that we observed are consistent with previous work in rats exposed to ethanol or THC during peri-adolescence. The effect of THC on reducing weight gain is consistent with recently published papers that used a similar THC exposure dose and route of administration (Bruijnzeel et al., 2019; Nelson et al., 2019a; Rubino et al., 2008; Scherma et al., 2016).

However, the increased sensitivity of females to the body weight effects of alcohol that we observed contrasts with effects reported in some studies (Amodeo et al., 2018; Fagundes et al., 2016; Forbes et al., 2013). This discrepancy may be due to differences in animal strain and alcohol consumption procedures in terms of onset, duration, route of administration, and dose. For example, the Amodeo et al (2018) study used intermittent alcohol vapor exposure, whereas Fagundes et al (2016) and Forbes et al (2013) modeled binge alcohol consumption (3 g/kg/day). The decrease in ethanol intake in the COM group compared to the EtOH group is also not entirely consistent with our previous study (Nelson et al., 2019a) where we observed THC-induced decreases in ethanol intake in rats that self-administered oral THC but not in those given s.c. injections of THC. One methodological change in the current study that may have contributed to these different findings was the shorter duration of drinking sessions during the dark cycle in our current work (1.5 h vs. 3 h in our previous study). There may be a unique interaction among THC, ethanol, and circadian rhythm (Friedman & Gershon, 1974), with THC administration inhibiting drinking activity or general activity level during the light portion of the daily cycle. The use of THC isolated from cannabis in the current study compared to the synthetic THC (dronabinol) used in Nelson et al. (2019a) may also be a factor contributing to these different results. Overall, these findings of body weight gain and ethanol intake implicate a prolonged metabolic impact of adolescent THC exposure in males compared to females, whose potential mechanisms require further investigation.

When brain slices were obtained from rats between 3 and 5 weeks after they ingested low to moderate levels of ethanol, were injected with THC, or were exposed to both drugs, we observed a reduced magnitude of HFS-induced LTD in the output layer (layers V/VI) of the mPFC. This effect was most robust in rats exposed to both drugs, with THC exposure alone having a more modest effect that was not statistically significant from controls. A previous study that investigated the impact of repeated exposure to the full agonist of CB_1_ receptors WIN 55,212-2 during early (P35-40) or mid-adolescence (P40-45) found the drug significantly impaired LTD in the mPFC that can be induced by HFS delivered in the ventral hippocampus (Cass et al., 2014). The less robust effect of THC that we observed here could be due to multiple factors, including partial activation of CB_1_ receptors by THC compared to the full activation by WIN 55212-2, or to differences in the abundance of cannabinoid receptors between the hippocampal-mPFC and intra-mPFC circuits (Marsicano & Kuner, 2008). Repeated exposure to THC with escalating doses (5-10 mg/kg/day) from P35 to P45 has been shown to reduce HFS-induced LTD in the PL region of the mPFC on P75 (Rubino et al., 2015), but recordings in this study were done in the presence of the GABA_A_ antagonist picrotoxin as a means to further decrease GABAergic tone. Future research should focus on the dynamics of co-localization of CB_1_ receptors and GABAergic interneurons in the mPFC during adolescence as an important contributing factor to the effects of repeated exposure to THC.

Combined exposure to ethanol and THC resulted in the largest reduction in HFS-induced LTD in deep layer mPFC neurons, which suggests an additive effect of ethanol and THC. This may be due to ethanol-induced potentiation of THC’s effects in the brain, which has previously been shown to include increased blood levels of THC and its metabolites (Hartman et al., 2015) and an extended half-life of THC (Toennes et al., 2011) when co-administered with ethanol.

Currently available literature on the interaction between alcohol and THC at synaptic levels is limited, but repeated co-use of the two drugs has been shown to reduce CB_1_ receptor-mediated miniature excitatory postsynaptic current frequency in Purkinje cells (Zou et al., 2022). In the PFC, however, inconsistent results have been reported for the effect of chronic ethanol consumption on CB_1_ receptor expression and binding affinity, as summarized in the review by Wolfe et al. (2022). To investigate potential mechanisms for the long-lasting disruption in HFS-induced plasticity that we observed in drug-exposed rats, we performed additional recordings in the presence of drugs that influence GABA signaling and activity in the GSK3β signaling pathway.

We found that perfusion of the GABA_A_ receptor positive allosteric modulator indiplon partially restored HFS-induced LTD in rats that drank ethanol and those that were exposed to both ethanol and THC. This suggests that exposure to these drugs during the stage of adolescence when the capacity to undergo HFS-induced plasticity in the mPFC is still developing (Caballero et al., 2014b; Kang et al., 2018) is due to a deficit in inhibitory tone in the mPFC. These findings are consistent with previous studies reporting impaired GABA maturation in binge-drinking rodents and human subjects. For instance, recordings of layer V pyramidal neurons in the PL of adult male rats exposed to vaporized ethanol for 14 h/day from P28 to P42 revealed a significant reduction in GABA_A_ receptor-mediated tonic currents that were hypothesized to be caused by long-term downregulation of GABA_A_ receptors (Centanni et al., 2017). Silveri et al. (2014) found that young adult binge drinkers have reduced GABA levels in the anterior cingulate cortex, a structure relevant to executive functioning. Another study also found long-lasting hypersensitivity of GABA_A_ receptors in the dentate gyrus to ethanol challenge after chronic intermittent ethanol exposure during adolescence (Fleming et al., 2012). On the other hand, although the exposure to THC alone during peri-adolescence did not significantly affect HFS-induced plasticity in early adulthood, perfusion of indiplon partially restored LTD in THC group compared to DMSO. This result suggests that the increase in HFS-induced fEPSP under DMSO condition in THC group may be due to impairment in GABA functioning. This is likely due to the fact that CB_1_ receptors, which THC is a partial agonist for, are found in abundance on cholecystokinin (CCK)-positive GABAergic interneurons in the frontal cortex (Marsicano & Kuner, 2008). Indeed, WIN 55212-2 (a full agonist for CB_1_ receptors) perfusion has been found to significantly reduce GABA release (Katona et al., 1999). Also important will be investigations of a potential role for type 2 cannabinoid (CB_2_) receptors in the effects of THC exposure on inhibition in the local mPFC circuit. THC is also a partial agonist at CB_2_ receptors and their stimulation has been shown to inhibit GABA_A_ receptor-mediated inhibitory postsynaptic response in Purkinje cells in the cerebellum (Sadanandan et al., 2020). CB_2_ receptors are found in pyramidal neurons in layers III and V of cerebral cortex (Onaivi, 2006), although their abundance and distribution in the mPFC has not been specified, nor has their presence on GABAergic neurons in the mPFC been investigated. Functionally, CB_2_ receptor activation ameliorated anxiety-like behaviors on the elevated plus maze in mice that underwent chronic alcohol exposure (four weeks of daily 4 g/kg alcohol via oral gavage; Li et al., 2023).

Therefore, there is a potential for the binding of THC on CB_2_ receptors to influence GABA signaling in mPFC circuit in the long run. Interestingly, the perfusion of AR-A014418, a GSK3β inhibitor, only rescued LTD in the combined-exposure (COM) group. Acute alcohol or THC exposure has been reported to increase phosphorylation of Akt and GSK3β in laboratory rodents (Neznanova et al., 2009; Ozaita et al., 2007). Chronic intake, on the other hand, may reduce phosphorylation of Akt and GSK3β (He et al., 2006). Because GSK3β phosphorylation (usually at the S9 site) inhibits its activity, chronic ethanol and THC exposure may lead to hyperactive GSKβ. Under this theoretical framework, AR-A014418 perfusion should compensate for the heightened GSK3β functioning and restore the HFS-induced LTD in ethanol- and THC-treated groups, not just the combined group. We hypothesize that the use of ethanol and THC alone may have influenced other systems or pathways in the downstream of Akt-GSK3β, and co-use of the drugs led to a “double hit” and detectable damage to these systems or pathways. For instance, GSK3β has been shown necessary to dopamine D_2_ receptor-mediated inhibition of NMDA receptors in layer V pyramidal neurons in PFC (Li et al., 2009). Since chronic intake of both ethanol and THC have been shown to reduce the CB1-dependent inhibition of glutamate signaling (Colizzi et al., 2016; Rao et al., 2015), the combined exposure could produce additive effects on the excitatory/inhibitory balance in mPFC, so that the local circuit is more sensitive to the perfusion of AR-A014418. Further research is required to elucidate the interaction of THC and ethanol on the Akt-GSK3β pathway, but a recent study by showed that knock-out of GSK3β in parvalbumin-positive (PV+) interneurons in the PFC increased their excitability and excitatory synaptic strength (Monaco et al. (2020). It may be that PV+ interneurons are an important component of the GSK3β-mediated reduction in LTD after long-term co-exposure to ethanol and THC.

A surprising finding in these studies was the apparent ability of DMSO to increase the fEPSP slope following HFS in brain slices from rats in the THC and COM groups. To account for this excitatory effect of DMSO, we adjusted fEPSP responses in indiplon and AR-A014418 conditions (see Supplementary Table 3 and Supplementary Figure 3), and the conclusion remained the same for the effects of indiplon and AR-A014418 in EtOH, THC, and COM groups.

This excitatory effect was unexpected in light of the widely accepted view that DMSO, under low concentrations, is a biologically inert solvent (Nasrallah et al., 2008). However, Tamagnini et al. (2014) reported that slice incubation concentrations of DMSO as low as 0.05% can increase the intrinsic excitability of mice hippocampal CA1 pyramidal cells by reducing membrane resistance and action potential threshold. An earlier study by Tsvyetlynska et al. (2005) showed that 0.02% DMSO can increase NMDA- and AMPA-mediated excitatory postsynaptic potential in lamprey reticulospinal neurons. In addition, a dose-dependent decrease in GABA-induced chloride current was reported in cultured rat dorsal root ganglion neurons exposed to 0.3-3% DMSO (Nakahiroa et al. (1992). Taken together, these findings suggest an increase in excitatory and decrease in inhibitory outputs under the influence of low-concentration DMSO, which may exacerbate disruptive effects of ethanol and THC on GABAergic signaling. The significant decrease in fEPSP slope with the perfusion of indiplon compared to the DMSO condition in the ethanol, THC, and combined treatment groups is consistent with this hypothesis, but additional studies are necessary to properly test it.

Functionally, the HFS-induced LTD in mPFC may contribute to the fine-tuning of deep layer pyramidal neural output that focuses its activity on task-relevant items instead of the background noise (Caballero et al., 2016; Kang et al., 2018; Seamans et al., 2001). Reduction of LTD caused by alcohol and THC co-use could potentially reflect disruption of adolescent-emerging inhibitory tone in mPFC mediated by GABA. Less GABAergic inhibitory neurotransmission is associated with poorer self-control and increased risky behaviors. Indeed, a study showed that moderate drinking can lead to increased inhibitory tone on dopaminergic neurons in ventral tegmental area, which implicates increased ratio of phasic-to-tonic firing pattern and increased DA release in response to risky stimulus (Spear, 2018). Endogenous endocannabinoid signaling in the PFC influences executive functioning; THC suppresses GABA release in the PFC possibly due to presynaptic CB_1_ receptors on GABAergic interneurons, which drives glutamatergic output from pyramidal neurons to the VTA, increasing VTA DA activities (Egerton et al., 2006). This delayed maturation of GABAergic system in mPFC caused by co-use of alcohol and cannabis may explain some cognitive and behavioral consequences in rodents, such as anxiety-like behaviors (Hamidullah et al., 2021), short-term memory deficit (Ciccocioppo et al., 2002; Hamidullah et al., 2021; Silva-Pena et al., 2019), and reversed novelty-seeking (Swartzwelder et al., 2012). Abnormal development of inhibitory tone in mPFC has been indicated in the psychopathology of many adolescent-onset diseases including schizophrenia (Renard et al., 2018) and drug addiction (Goldstein & Volkow, 2011). Therefore, co-use of THC and ethanol could potentially render adolescents more vulnerable to develop these diseases. Indeed, clinical research has associated THC and alcohol co-use with higher risk of alcohol use disorder (Hayaki et al., 2016; Liang, 2022; Subbaraman & Kerr, 2015; Thompson et al., 2021; Waddell, 2021), poorer selective attention accuracy (Jacobus et al., 2015; Ramaekers et al., 2011; Wade et al., 2020a), greater discounting of future rewards (Petker et al., 2021), poor learning, memory, and visuospatial functioning (Bedillion et al., 2021). Our study suggests interventions that target GABA signaling may be a promising target for pharmacological interventions aimed at addressing deficits associated with the co-use of alcohol and cannabis, as well as pointing to the unique impairment of Akt-GSK3β pathway that was not caused by use of either drug alone.

## ACKNOWLEDGEMENTS

The authors thank Anna Tsyrulnikov, Avni Dave, Michael Wostmann, Vincent Marsh, and Rashmi Ghonasgi for their assistance in drug administration part of the experiment. The authors also thank Holly Fairfield for animal care and colony maintenance. This work was supported in part by funding from The National Institute on Drug Abuse (R21 DA 045175) awarded to NCL and JMG.

