## Supplementary Material for "Effects of combined exposure to ethanol and delta-9-tetrahydrocannabinol during adolescence on synaptic plasticity in the prefrontal cortex of Long Evans rats"

**Supplementary Table 1**

**Treatment Group Assignment**. Rats were pseudorandomly assigned to groups so that intake of saccharin during the habituation sessions on P28-P29 was approximately equal. Shown in the table are the individual rat IDs, their average body weight over the two-day habituation, and consumption of the 0.1% saccharin (SACC) solution on each day’s 2.5h session.

CTL – control; EtOH – ethanol only; THC – THC only; COM – ethanol and THC

| **THC** | **Mean Body Weight (g)** | **SACC**  **Day 1 (mL)** | **SACC**  **Day 2 (mL)** |  | **CTL** | **Mean Body Weight (g)** | **SACC**  **Day 1 (mL)** | **SACC**  **Day 2 (mL)** |
| --- | --- | --- | --- | --- | --- | --- | --- | --- |
| GL2-3 | 48.6 | 1.8 | 2.5 |  | GL2-5 | 41.5 | 1.1 | 0.9 |
| GL2-4 | 44.5 | 1.7 | 2.9 |  | GL2-6 | 44.3 | 2.6 | 4.4 |
| GL2-21 | 58.1 | 5.8 | 5.1 |  | GL2-19 | 59.4 | 4.2 | 2.4 |
| GL2-22 | 54.9 | 3.2 | 1.4 |  | GL2-20 | 66.2 | 3.4 | 3.4 |
| GL2-33 | 54.4 | 1.7 | 4.3 |  | GL2-37 | 52.5 | 2.5 | 3.1 |
| GL2-34 | 53.1 | 4.2 | 2.7 |  | GL2-38 | 50.9 | 4.2 | 3.4 |
| GL2-49 | 75.8 | 3.3 | 5.2 |  | GL2-55 | 77.1 | 4.1 | 9 |
| GL2-50 | 74.4 | 5.3 | 6.7 |  | GL2-56 | 76.4 | 2.5 | 2.6 |
| Mean | 58.0 | 3.4 | 3.9 |  | Mean | 58.5 | 3.1 | 3.7 |
| SD | 11.4 | 1.6 | 1.8 |  | SD | 13.7 | 1.1 | 2.4 |
| **COM** | **Mean Body Weight (g)** | **SACC**  **Day 1 (mL)** | **SACC**  **Day 2 (mL)** |  | **EtOH** | **Mean Body Weight (g)** | **SACC**  **Day 1 (mL)** | **SACC**  **Day 2 (mL)** |
| GL2-7 | 47.4 | 1.6 | 3.2 |  | GL2-1 | 40.7 | 5.5 | 5.4 |
| GL2-8 | 47.0 | 2.3 | 4.7 |  | GL2-2 | 41.0 | 2.9 | 3.7 |
| GL2-23 | 58.2 | 4.5 | 3.9 |  | GL2-17 | 55.6 | 3.6 | 3.5 |
| GL2-24 | 55.6 | 2.9 | 3.2 |  | GL2-18 | 58.9 | 3.3 | 3.5 |
| GL2-39 | 49.9 | 3.1 | 3.2 |  | GL2-35 | 48.5 | 3.3 | 3.6 |
| GL2-40 | 51.2 | 3.7 | 2.9 |  | GL2-36 | 49.6 | 3.7 | 4.2 |
| GL2-51 | 80.3 | 1.4 | 6.9 |  | GL2-52 | 80.6 | 6.5 | 6.7 |
| GL2-53 | 73.5 | 6.0 | 8.0 |  | GL2-54 | 74.3 | 4.0 | 5.7 |
| Mean | 57.9 | 3.2 | 4.5 |  | Mean | 56.1 | 4.1 | 4.5 |
| SD | 12.5 | 1.5 | 1.9 |  | SD | 14.7 | 1.2 | 1.2 |

**Supplementary Figure 1**

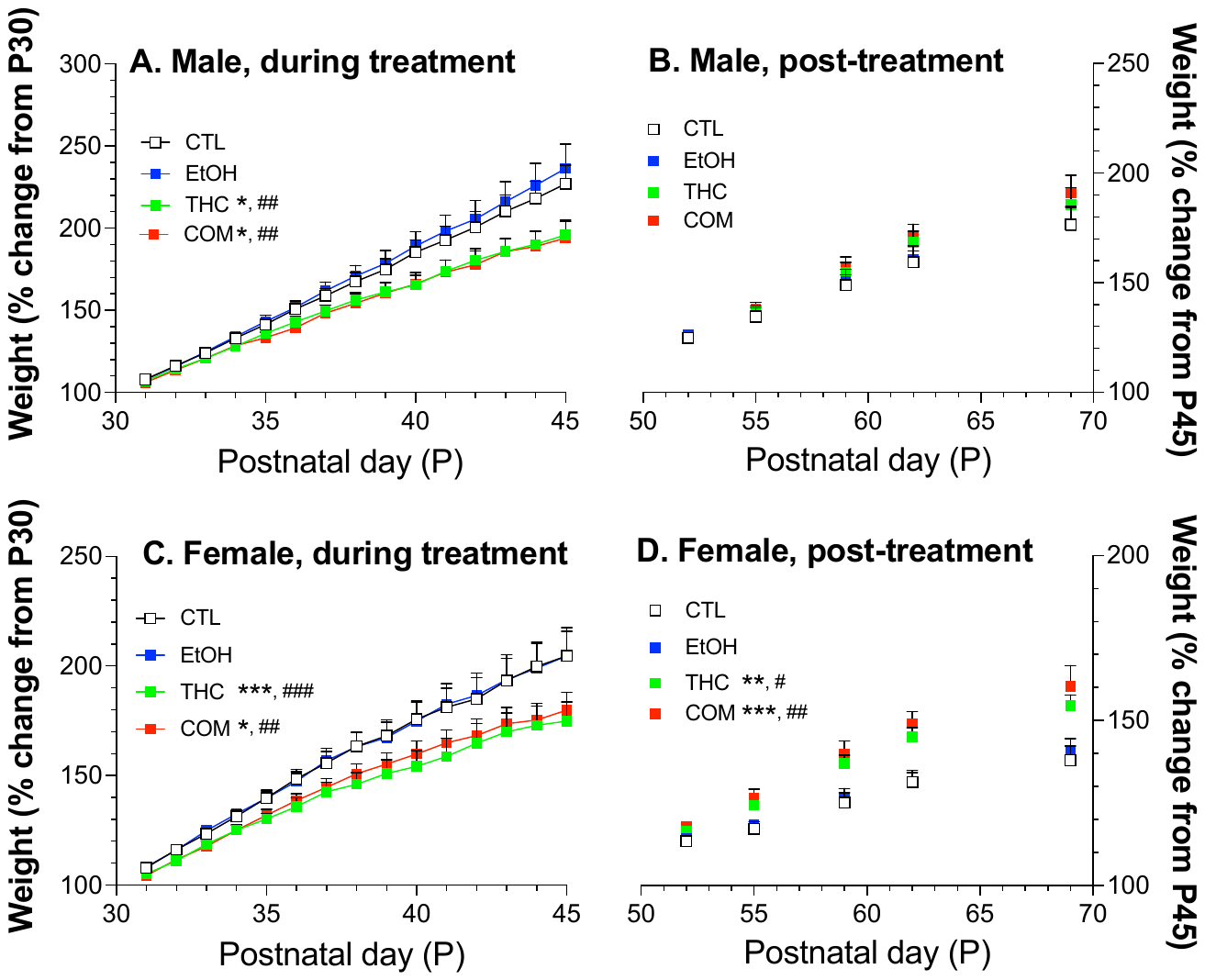

Supp. Fig. 1. Effects of ethanol, THC, or their combination on growth rate of males and females. Weights were measured daily P24-P45, and on P52, P55, P59, P62, and P69 (n = 7-8 rats/sex/group). Daily weight gains during drug exposure (P31-45; panels A and C) and following treatment (P52-P69; panels B and D) were analyzed for each sex using two-way ANOVA (day x treatment). Separate analyses were performed for weight during drug exposure (P31-P45, measured daily) and after drug exposure (P52-69, measured on P52, 55, 59, 62, and 69). Main effects of treatment (males: F_3, 405_ = 28.0, *p* < 0.001; females F_3, 420_ = 24.5, *p* < 0.001) and day (males: F_14, 405_ = 100, *p* < 0.001; females: F_14, 420_ = 69.5, *p* < 0.001) were found during treatment period. No significant interactions were found. Post-hoc analysis revealed that THC-treated groups (THC and COM) had significantly reduced body weight gain compared to CTL and EtOH groups (panels A and C) for both sexes. During the post-treatment period (P52-69), there was only a significant effect of treatment in females (F_3, 140_ = 24.9, *p* < 0.001) but not males. Day of the experiment still had a main effect in both sexes (males: F_4, 135_ = 74.8, *p* < 0.001; females: F_4, 140_ = 76.6, *p* < 0.001). No significant interaction between day and treatment was found. Post-hoc analysis showed that THC and COM groups had significantly reduced body weight gain compared to CTL and EtOH in females (panel D). **p* < 0.05, ***p* < 0.01, and ****p* < 0.001, compared to CTL; ^#^*p* < 0.5, ^##^*p* < 0.01, and ^###^*p* < 0.001 compared to EtOH. CTL – control; EtOH – ethanol only; THC – THC only; COM – ethanol and THC

**Supplementary Figure 2**

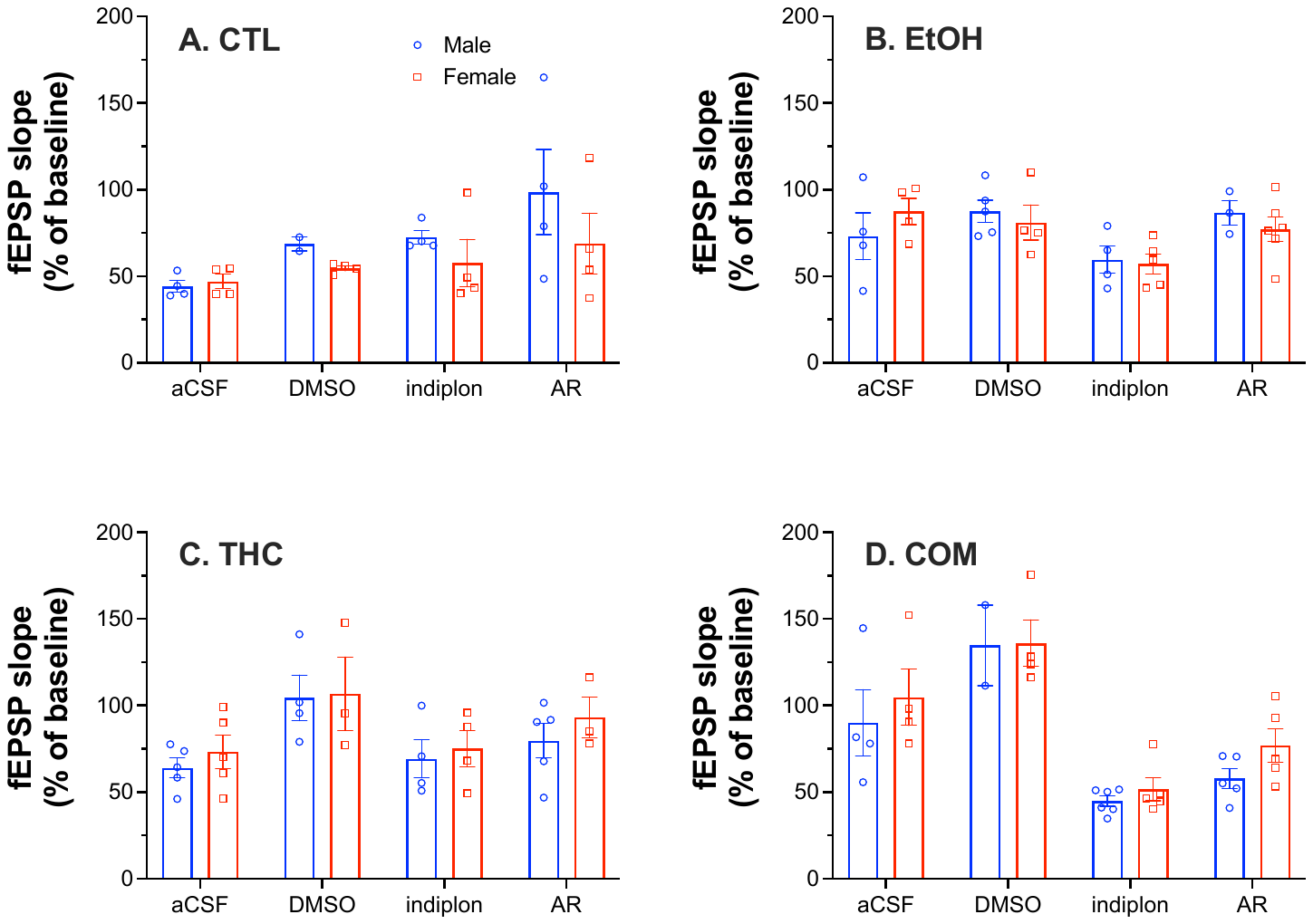

Supp. Fig. 2. Effects of 0.1-0.2% DMSO, 5 μM indiplon, and 10 μM AR-A014418 on HFS-induced changes in the slope of the fEPSP in brain slices from rats in CTL, EtOH, THC, and COM group respectively. Plotted are the mean fEPSP slopes for the 30-40 min period post-HFS to represent long-term stable response to HFS. Data split by sex for presentation purposes only. *n* = 6-11 slices from 6-11 rats/group.

**Supplementary Table 2**

**Average percent change in the fEPSP slope 30-40 min post-HFS, normalized to the 10-min baseline period pre-HFS.** Slices were recorded for 10 min in aCSF only to establish a baseline, and an additional 10 min baseline for DMSO, IND, and AR conditions with the respective perfusate (DMSO: DMSO only; IND – Indiplon + DMSO; AR – AR-A014418 + DMSO). HFS was then delivered (4 trains of 100 pulses at 50 Hz/train; 10-s inter-train interval), followed by a 40-min recording period throughout which the perfusion continued. All values are in the form of mean ± SEM. CTL – control; EtOH – ethanol only; THC – THC only; COM – ethanol and THC. **p* < 0.05, ***p* < 0.01, and ****p* < 0.001 vs. aCSF; **^#^***p* < 0.05, **^##^***p* < 0.01, and **^###^***p* < 0.001 vs. DMSO.

|  | CTL | | EtOH | | THC | | COM | |
| --- | --- | --- | --- | --- | --- | --- | --- | --- |
|  | Mean  (± SEM) | Sample Size  (M, F) | Mean  (± SEM) | Sample Size  (M, F) | Mean  (± SEM) | Sample Size  (M, F) | Mean  (± SEM) | Sample Size (M, F) |
| aCSF | 45.5  (2.52) | 8  (4, 4) | 80.2  (7.66) | 8  (4, 4) | 68.7  (5.47) | 10  (5, 5) | 97.5  (12.0) | 8  (4, 4) |
| DMSO | 59.2  (3.25) | 6  (2, 4) | 84.7**^#^**  (5.47) | 9  (5, 4) | 106** (10.7) | 7  (4, 3) | 136* (10.4) | 6  (2, 4) |
| IND | 64.6  (7.11) | 8  (4, 4) | 58.2  (4.46) | 9  (4, 5) | 72.2**^#^**  (7.14) | 8  (4, 4) | 47.9***^,^**^#^** (3.39) | 11  (6, 5) |
| AR | 83.7***  (15.1) | 8  (4, 4) | 80.3  (5.30) | 9  (3, 6) | 84.8  (7.47) | 8  (5, 3) | 67.4*^,^ **^###^** (6.16) | 10  (5, 5) |

**Supplementary Table 3**

**Mean percent change in the fEPSP slope post-HFS, adjusted to DMSO.** DMSO (0.1%-0.2%) was used as the solvent to dissolve indiplon and AR-A014418. To account for the changes in the fEPSP response that was associated with DMSO perfusion, the mean % fEPSP slopes from slices perfused with indiplon and AR-A014418 were adjusted to the DMSO/aCSF value. DMSO/aCSF value was calculated by dividing the mean % fEPSP slope during the DMSO condition of a treatment group (e.g., 59.2 for CTL DMSO) by the mean % fEPSP slope of the aCSF condition from the same group (e.g., 45.5 for CTL aCSF). This reflects the ratio of the increase in fEPSP response with DMSO perfusion. The adjusted mean was calculated by dividing the initial mean (which are shown in Table 2) by the corresponding value of DMSO/aCSF. One-way ANOVA for each treatment group with three conditions (aCSF, indiplon, AR-A014418) revealed significant effects of perfusion condition in EtOH (F_2, 23_ = 5.66, *p* = 0.010), THC (F_2, 23_ = 4.78, *p* = 0.018), and COM (F_2, 26_ = 24.2, *p* < 0.001). Post-hoc analyses: **p* < 0.05 and ****p* < 0.001 vs. aCSF within each treatment group. CTL – control; EtOH – ethanol only; THC – THC only; COM – ethanol and THC.

|  | indiplon | | | AR-A014418 | | |
| --- | --- | --- | --- | --- | --- | --- |
|  | DMSO  /aCSF | Initial Mean  (± SEM) | Adjusted Mean  (± SEM) | DMSO  /aCSF | Initial Mean  (± SEM) | Adjusted Mean  (± SEM) |
| CTL | 1.30 | 64.6  (7.11) | 49.7  (5.47) | 1.30 | 83.7  (15.1) | 64.4  (11.6) |
| EtOH | 1.06 | 58.2  (4.46) | 55.2* (4.23) | 1.06 | 80.3  (5.30) | 76.0  (5.02) |
| THC | 1.54 | 72.2  (7.14) | 47.0*  (4.65) | 1.54 | 84.8  (7.47) | 55.2  (4.86) |
| COM | 1.39 | 47.9  (3.39) | 34.4***  (2.44) | 1.39 | 67.4  (6.16) | 48.4***  (4.42) |

**Supplementary Figure 3**

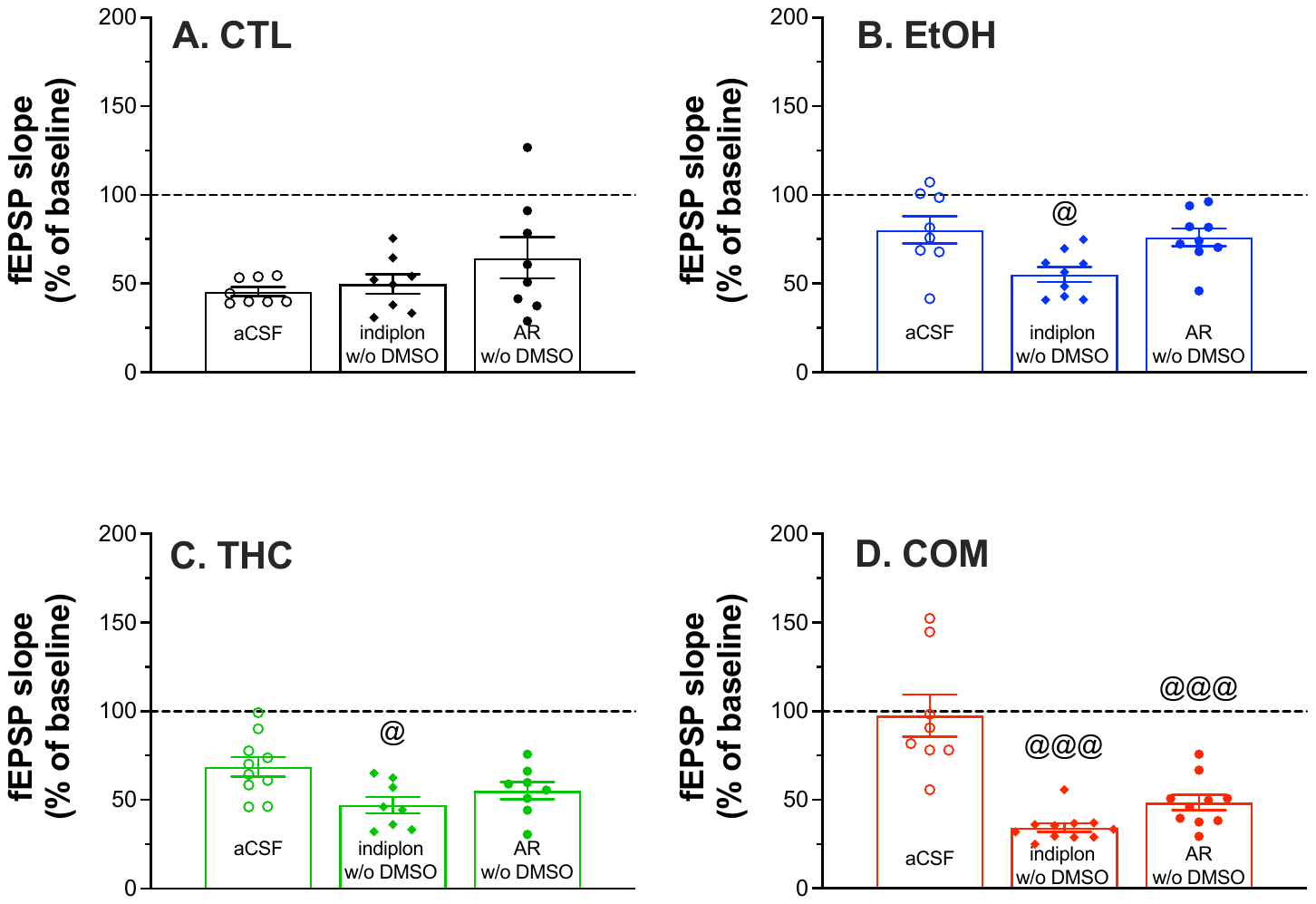

Suppl. Fig. 3. Effects of 0.1-0.2% DMSO, 5 μM indiplon, and 10 μM AR-A014418 on HFS-induced changes in the slope of the fEPSP in brain slices from rats in CTL, EtOH, THC, and COM group respectively. Plotted are the mean fEPSP slopes for the 30-40 min period post-HFS to represent long-term stable response to HFS. Datapoints in indiplon and AR groups have been adjusted to the mean response in DMSO condition such that each value was divided by the average fold increase in normalized aCSF slope from aCSF to DMSO (see Table 3 for the calculation). ^@^*p* < 0.05 and ^@@@^*p* < 0.001 vs. aCSF. *n* = 6-11 slices from 6-11 rats/group.
